# 1D model simulations suggest biofilms can suppress cathodic H_2_ bubble formation

**DOI:** 10.64898/2026.11.03.763859

**Authors:** Mikkel Bendix Jensen, Vladimir Koselka, Jacopo Catalano, Jo Philips

**Author notes:** These authors contributed equally to this work (Mikkel Bendix Jensen, Vladimir Koselka).

## Abstract

Microbial electrosynthesis (MES) electrochemically generates H_2_ for the reduction of CO_2_ into acetate by acetogens. So far, the importance of a biofilm on the cathode of such systems remains under debate. Here, we investigated the role of cathodic biofilms through computational modelling. We developed MES-BIOFILM, a comprehensive one-dimensional model to simulate steady-state H_2_ and CO_2_ concentration gradients within cathodic biofilms, while accounting for the limited solubility of H_2_. The effects of key biofilm properties, including thickness, cell density and H_2_ consumption rate were analysed. Overall, our simulations indicate that cathodic biofilms play an important role in (1) minimizing H_2_ loss from the system through H_2_ consumption within the biofilm; and (2) suppressing H_2_ bubble formation by maintaining the H_2_ concentrations at the cathode surface below saturation. Our model predicts that the current density at which H_2_ bubbles begin to form can be increased by 5.5-fold relative to a bare electrode when biofilms exhibit a high cell density and high H_2_ consumption rate. This effect requires biofilms of at least 50 µm thickness, whereas CO_2_ diffusion limitations arise above 200 µm. These findings highlight the critical role of biofilms to increase MES performance, while requiring further experimental validation.

## Introduction

Microbial electrosynthesis (MES) is a promising biotechnological process for the conversion of CO_2_ into small organic molecules (e.g. acetate) (Nevin *et al*., 2010, Rabaey & Rozendal, 2010) by acetogenic bacteria according to the reaction (Drake *et al*., 2008):

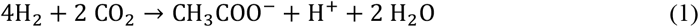

During MES, the required molecular hydrogen (H_2_) is electrochemically generated by a cathode placed in the bioreactor. This *in situ* H_2_ generation is advantageous in comparison to gas fermentation (i.e. feeding gaseous H_2_), as dissolving gaseous H_2_ is impeded by its low aqueous solubility (Table 1) (Carvajalino-Olave *et al*., 2025). Nevertheless, MES struggles with insufficient product formation rates, hindering its upscaling and industrial implementation (Prevoteau *et al*., 2020, Jourdin & Burdyny, 2021, Wood *et al*., 2021). One of the main issues limiting MES rates is the formation of H_2_ bubbles on the cathode when applying increased current densities, eliminating the advantage of MES versus gas fermentation (Kracke *et al*., 2021). In addition, the loss of H_2_ to the headspace lowers the efficiency of MES systems (LaBelle & May, 2017, Ragab *et al*., 2020, Kracke *et al*., 2021, Kuchenbuch *et al*., 2025).

**Table 1:** Parameter values used for MES-BIOFILM model.

| Parameter | Symbol | Value | Unit | Reference |
|---|---|---|---|---|
| Faraday’s constant | *F* | 96485.3321 | s · A · mol<sup>−1</sup> | (Newell & Tiesinga, 2019) |
| Diffusion coefficient of H<sub>2</sub> in water at 25°C | *D*<sub>H<sub>2</sub>,water</sub> | 4.5 · 10<sup>−9</sup> | m<sup>2</sup> · s<sup>−1</sup> | (Cussler, 2009) |
| Diffusion coefficient of CO<sub>2</sub> in water at 25°C | *D*<sub>DIC,water</sub> | 1.92 · 10<sup>−9</sup> | m<sup>2</sup> · s<sup>−1</sup> | (Cussler, 2009) |
| Diffusion coefficient of H<sub>2</sub> in biofilm | *D*<sub>H<sub>2</sub>,bf</sub> | 2.27 · 10<sup>−9</sup> | m<sup>2</sup> · s<sup>−1</sup> | Determined in this study (Fig S1) |
| Diffusion coefficient of DIC in biofilm | *D*<sub>DIC,bf</sub> | 0.562 · 10<sup>−9</sup> | m<sup>2</sup> · s<sup>−1</sup> | Estimated in this study (Fig S1) |
| Saturation concentration of H<sub>2</sub> in water at 30°C | *C*<sub>H<sub>2</sub>,sat</sub> | 802 · 10<sup>−3</sup> | mol · m<sup>−3</sup> | (Kolev, 2011) |
| Saturation DIC concentration in water at pH 7 | *C*<sub>DIC,sat</sub> | 227 | mol · m<sup>−3</sup> | Calculated using Visual Minteq as the DIC concentration in water at pH 7 in equilibrium with a 100% CO<sub>2</sub> headspace at 1 atm |
| Thickness diffusive boundary layer | λ | 200 | µm | (Rivera-Alvarez *et al.*, 2020) |
| Mass transfer coefficient of H<sub>2</sub> | *h*<sub>H<sub>2</sub></sub> | 22.5 | µm · s<sup>−1</sup> | Calculated in this study (Eq 22) |
| Mass transfer coefficient of DIC | *h*<sub>DIC</sub> | 9.6 | µm · s<sup>−1</sup> | Calculated in this study (Eq 22q) |
| Biofilm thickness | *L* | 10 − 1000 | µm | Chosen for this study |
| Biofilm cell density | *C*<sub>*x*</sub> | 10 − 400 | gDW · L<sup>−1</sup> | Table S1 |
| Threshold concentration of H<sub>2</sub> | H<sub>2</sub><sup>⋆</sup> | 0 − 4.4 · 10<sup>−5</sup> | mol · m<sup>−3</sup> | (Munoz & Philips, 2023) |
| Biomass-specific first order rate coefficient for H<sub>2</sub> consumption | *k*<sub>1</sub><sup>\*</sup> | 10 − 60 | (h · gdW · L<sup>−1</sup>)<sup>−1</sup> | (Munoz-Duarte *et al.*, 2025) |
| Monod half saturation constant for DIC | *K*<sub>*m*</sub> | 4 · 10<sup>−4</sup> | mol · m<sup>−3</sup> | (Callander *et al.*, 2024) |

Several scientists have suggested that a cathodic biofilm, i.e. a layer of microbial cells attached to the cathode surface by a matrix of extracellular polymeric substances (EPS), is crucial for enhancing the productivity of MES (Jourdin & Burdyny, 2021, Vassilev *et al*., 2022). Microbial cells attached to the cathode surface could indeed consume the H_2_ generated by the cathode, before it diffuses to the bulk liquid and ultimately escapes to the headspace. So far, however, cathodic biofilms are considered too thin, especially if using pure cultures of acetogens (comprehensive overview given in (Grøn *et al*., 2026)). One study successfully increased MES rates by embedding the acetogen *Sporomusa ovata* in a thick hydrogel-based synthetic biofilm (1 mm) (Krige *et al*., 2021). However, biofilms could possibly also be too thick, as a thick layer would restrict the access to CO_2_ due to diffusion limitations (Vassilev *et al*., 2022). So far, the optimal thickness of a cathodic biofilm has not been determined. Even the necessity of cathodic biofilms for MES remains unclear, as some studies have obtained good MES rates with purely planktonic cultures (Deutzmann & Spormann, 2024).

One possibly beneficial effect of cathodic biofilms that has not yet been considered is the suppression of H_2_ bubble formation. Increasing cathodic current densities leads to H_2_ bubble formation once the dissolved H_2_ concentration on the cathode surface reaches or surpasses its solubility limit (H_2_ saturation) (Carvajalino-Olave *et al*., 2025). H_2_ consumption in a cathodic biofilm could keep the H_2_ concentration at the cathode surface below saturation, thereby preventing H_2_ bubble formation. However, at too high current densities, a biofilm would not be capable anymore of maintaining the H_2_ concentration below saturation, leading to the formation of H_2_ bubbles, ultimately disrupting the biofilm (Figure 1) (Rominger *et al*., 2025). So far, the extent to which cathodic biofilms can suppress H_2_ bubble formation upon increasing cathodic current densities has not been investigated.

**Figure 1:**
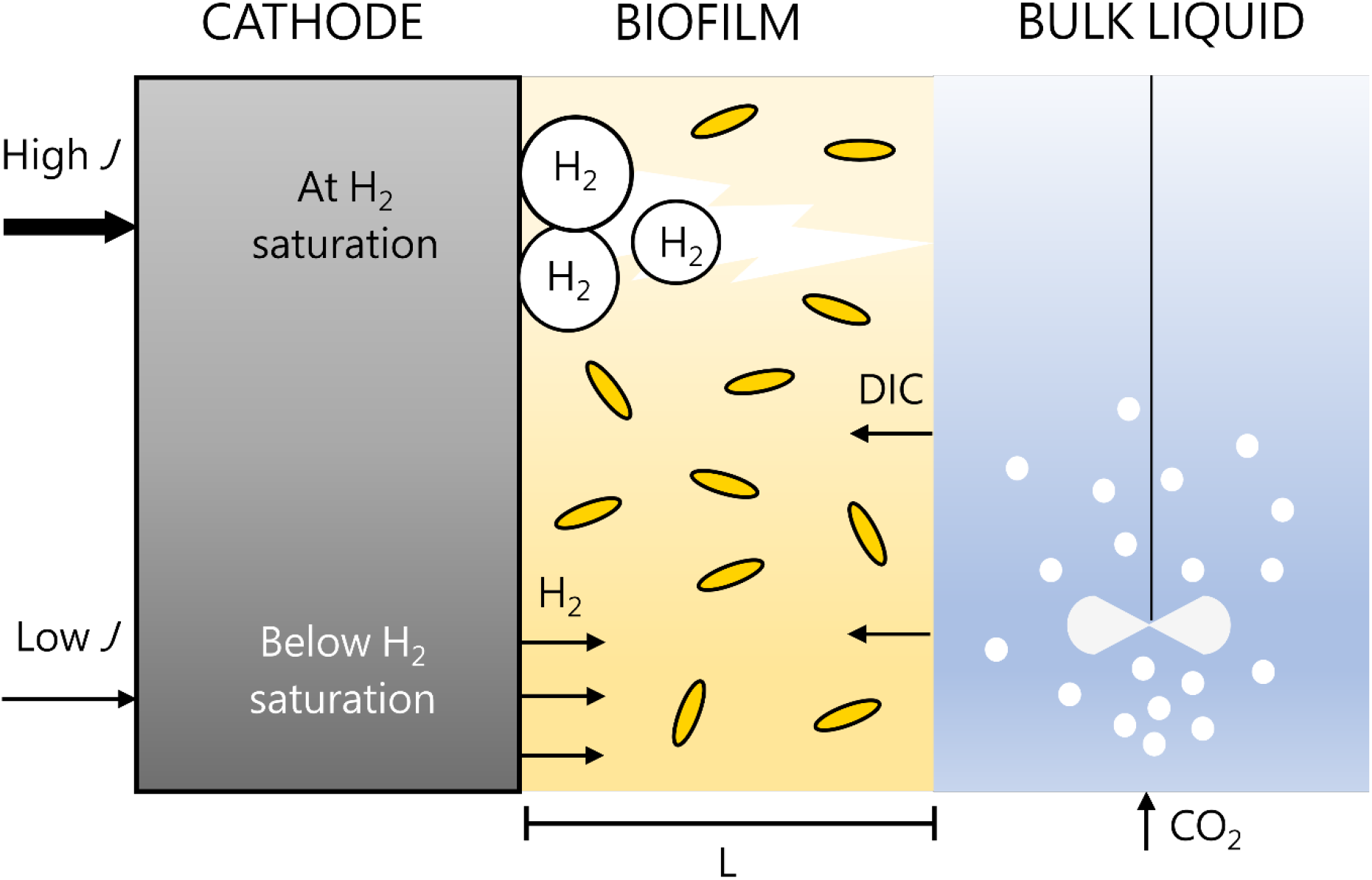
Scheme illustrating the biofilm described by the model MES-BIOFILM. The biofilm is a one-dimensional layer with thickness L and with homogeneously spread acetogen cells, which consume H_2_ and CO_2_. The biofilm is exposed on the right side to a bulk liquid, which is sparged with CO_2_ gas and thus provides DIC to the biofilm. The biofilm is attached to a cathode. When low current densities are supplied to the cathode, the generated H_2_ remains dissolved and diffuses into the biofilm. When the current density is so high that the saturated H_2_ concentration is reached at the cathode surface, H_2_ bubbles are formed and disrupt the biofilm. It the latter case, MES-BIOFILM is not applicable anymore.

Insights into H_2_ and CO_2_ concentration gradients in cathodic biofilms would be highly valuable to assess optimal biofilm thickness, as well as the possibility of H_2_ bubble suppression. Unfortunately, such concentration gradients are difficult to measure, definitely in thin cathodic biofilms (Sandfeld *et al*., 2024). Alternatively, computational modelling offers an interesting tool to predict concentration gradients. Various models have already been developed to gain insights into the factors limiting MES rates, as well as the role of biofilms in bioelectrochemical systems (review of different models given by (Li *et al*., 2022)). Cabau-Peinado et al. (2021) already modelled how MES operational parameters affect the growth of a cathodic biofilm. Kazemi *et al*. (2015) simulated concentration gradients in cathodic biofilms, but assumed a direct extracellular electron transfer (i.e. no H_2_). Recently, Murugesan et al. (2026) modelled both H_2_ and CO_2_ concentration gradients in cathodic biofilms, but did not consider the limited H_2_ solubility nor H_2_ gas evolution.

In this work, we developed a comprehensive 1D computational model, called MES-BIOFILM, to simulate H_2_ and CO_2_ concentration gradients in steady-state cathodic biofilms of acetogens. Our model incorporated diffusion and microbial consumption of H_2_ and CO_2_ in the biofilms. In addition, MES-BIOFILM accounted for the limited solubility of H_2_ by setting H_2_ saturation as the maximum feasible H_2_ concentration. We first simulated thin biofilms (up to 200 µm), in which the CO_2_ concentration is likely never a limiting factor with realistic current densities. We modelled the effect of various biofilm properties (i.e. biofilm thickness, cell density, H_2_ consumption rate) on the H_2_ concentration gradients and evaluated which properties allow for 100% conversion efficiency (no loss of H_2_ to bulk liquid). In addition, we determined to which extent biofilms can increase the current density at which H_2_ saturation gets reached. Next, we simulated thick biofilms (up to 1 mm) and evaluated at which biofilm thicknesses CO_2_ limitations arise. Overall, our simulation results provide insights into the role of cathodic biofilms in suppressing H_2_ bubble formation and can thereby help guide the design of effective MES systems.

## Methods

### Model concept and assumptions

In this study, the comprehensive model MES-BIOFILM was developed to describe the diffusion and microbial consumption of H_2_ and CO_2_ in cathodic biofilms. This model describes the biofilm as a one-dimensional layer with thickness *L* [m], in which acetogen cells are homogenously distributed with a cell density *C*_x_ [g_DW_·L^-1^] (Figure 1). This biofilm is considered attached to the cathode at one side and exposed on the other side to a well-mixed aqueous bulk liquid, which removes dissolved H_2_ and provides CO_2_ through convection over the biofilm surface. The CO_2_ dissolves as CO_2,aq_, HCO_3_^-^ and CO_3_^2-^ (together dissolved inorganic carbon (DIC)), which diffuse through the biofilm. As acetogens have carbonic anhydrase enzymes to reversibly convert HCO_3_^-^ into CO_2_ (Braus-Stromeyer *et al*., 1997) and we assumed a fixed pH (7), MES-MODEL considered the lumped DIC concentration as the form of CO_2_ utilized by the microbes, without resolving the individual contribution of the different dissolved CO_2_ species.

MES-BIOFILM further assumes that a certain current density *J* (A·cm^-2^) is applied to the cathode, leading to electrochemical H_2_ evolution with 100% Faradaic efficiency (no side reactions). In addition, it was assumed that the H_2_ generated by the cathode remains dissolved and diffuses through the biofilm, as long as the dissolved H_2_ concentration remains below H_2_ saturation (Figure 1, upper case). However, once H_2_ saturation is reached, we assume that H_2_ bubble formation occurs, disintegrating the biofilm and turning our model simulations invalid (Figure 1, lower case).

MES-BIOFILM only considers steady-state conditions. Our model includes H_2_ and DIC consumption by the acetogenic bacteria but does not simulate pH gradients and assumes that pH and the acetate concentration were not affecting the rate of H_2_ and DIC consumption.

### Mathematical description

#### Dissolved H_2_ concentration

The change of the dissolved H_2_ concentration in the biofilm can be described by one-dimensional diffusion of H_2_ through the biofilm and the microbial consumption of H_2_ in the biofilm, given by the partial differential equation:

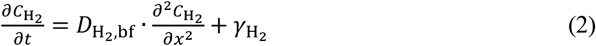

Where 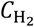 is the local dissolved H_2_ concentration in the biofilm [mol·m^-3^·s^-1^] at a distance *x* [m] from the cathode and at time *t* [s], 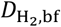 is the binary diffusion coefficient of H_2_ in the biofilm [m^2^·s^-1^], and *γ*_H_ is the microbial H_2_ consumption rate [mol·m^-3^·s^-1^]. As steady-state conditions are assumed for the concentrations within the biofilm, Equation (2) is reduced to the ordinary second-order differential equation:

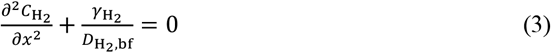

The microbial H_2_ consumption rate 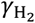 further depends on the dissolved H_2_ concentration following first-order kinetics as described for acetogens (Munoz-Duarte *et al*., 2025), while Monod kinetics was assumed for the dependence on the DIC concentration:

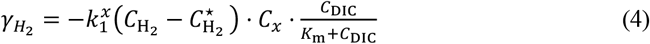

Here 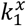 is the biomass specific first-order rate coefficient for the consumption of H_2_ by acetogens in [s^-1^·g_DW_^-1^·L] (Munoz-Duarte *et al*., 2025), 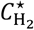 is the H_2_ threshold concentration given in [mol·m^-3^], which is the H_2_ concentration below which the H_2_ consumption halts (Munoz & Philips, 2023), *C*_x_ is the biofilm cell density in [g_DW_·L^-1^], *C*_DIC_ is the local DIC concentration inside the biofilm in [mol·m^-3^], and *K*_m_ is the half-saturation concentration for DIC in [mol·m^-3^]. In the case that CO_2_ was assumed not to be limiting, the factor 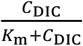 was then set to 1.

As left-hand boundary condition, it was assumed that the current density *J* applied to the cathode corresponds to constant H_2_ generation and hence a constant flux of H_2_ diffusing into the biofilm:

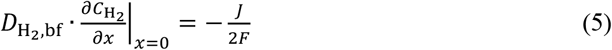

Where *J* is the current density provided to the cathode [A·m^-2^] and *F* is the Faraday constant [s·A·mol^-1^]. The factor 2 accounts for the two electrons required from the formation of one H_2_.

As right-hand boundary condition, it was assumed that the diffusive flux of H_2_ out of the biofilm equals the convective flux from the biofilm into the bulk liquid:

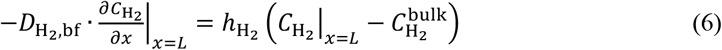

Where 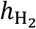 is the mass transfer coefficient of H_2_ [m·s^-1^] and 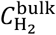 is the dissolved H_2_ concentration in the bulk liquid [mol·m^-3^], which was assumed to be zero.

#### DIC concentration

Similarly to the change in the dissolved H_2_ concentration, the change in the DIC concentration in the biofilm depends on diffusion of DIC through the biofilm and the microbial consumption of DIC in the biofilm, which in steady-state conditions gives the second-order ordinary differential equation:

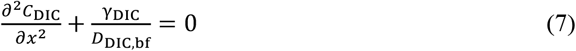

Where *C*_DIC_ is the local DIC concentration in the biofilm [mol·m^-3^], and *D*_DIC,bf_ is the binary diffusion coefficient of CO_2_ in the biofilm in [m^2^·s^-1^], assuming that CO_2_ transport can be used as a proxy for DIC transport. We further assumed that the microbial DIC consumption rate *γ*_DIC_ [mol·m^-3^·s^-1^] is half of the H_2_ consumption rate (Equation 4), because of the stoichiometry of Reaction 1:

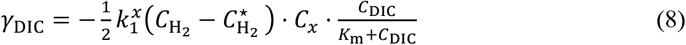

As left-hand boundary condition, no-flux was set, because of the impermeable (solid) cathode:

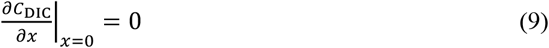

As the right-hand boundary condition, the diffusive flux of DIC into the biofilm was set equal to the convective flux of DIC from the bulk liquid into the biofilm:

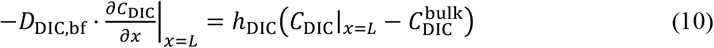

Where *h*_DIC_ is the DIC mass transfer coefficient [m·s^-1^] and 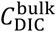 is the concentration of DIC in the bulk liquid [mol·m^-3^], which was assumed to be the DIC concentration in water at pH 7 in equilibrium with a headspace of 100% CO_2_ at 1 atm (Table 1). For the specific conditions of our simulations, the right-hand boundary condition can be altered into a fixed concentration boundary condition as derived in Supplementary Method 1:

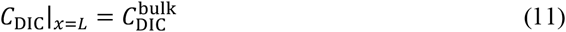

### Implementation in Python

The system of ODEs was solved in Python 3.12 using the SciPy’s solve_ivp function. The governing second-order ODEs (Equations 3 and 7) were first rewritten as for coupled first order ODEs:

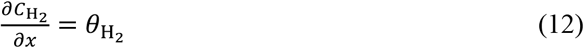

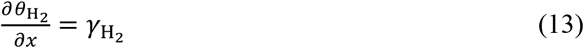

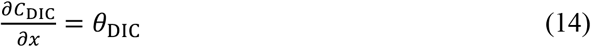

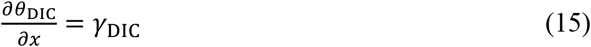

The resulting system of first-order ODEs was integrated from the left-hand boundary (at *x* = 0) to the right-hand boundary (*x* = *L*). The left-hand boundary conditions for Equation (12) and (14) were determined from the prescribed fluxes at *x* = 0, as specified by Equation (5) and (9), while the left-hand boundary conditions for Equation (13) and (15) were initially guessed. Next, an iterative minimization procedure was used to adjust the guessed concentration values until the resulting concentration profiles satisfied the two boundary conditions at the right-hand boundary within an acceptable error. For H_2_, the boundary condition at *x* = *L* prescribes the concentration gradient (here noted as 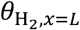) to be proportional to the convective flux (Equation 6). The corresponding error was calculated as the difference between the gradient obtained by the integrated solution and the prescribed value, both normalized by the prescribed gradient. For DIC, the right-hand boundary condition is a fixed DIC concentration (Equation 11). The corresponding normalized error was thus calculated from the difference between the obtained DIC concentration at *x* = *L* and the bulk concentration. The two errors were combined into a single objective function:

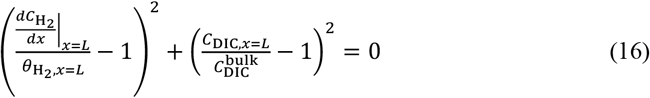

The minimization algorithm iteratively adjusted the initial guesses for the H_2_ and DIC concentrations until the objective function fell below 10^−6^, indicating that both right-hand boundary conditions were satisfied within the required numerical tolerance.

### H_2_ utilization efficiency of biofilm

The H_2_ utilization efficiency (*E*) of the biofilm is here defined as the fraction of the H_2_ generated by the cathode that gets converted within the biofilm. This can be calculated from the difference between the H_2_ flux at the left (Eq. 5) and right boundary of the biofilm, according to:

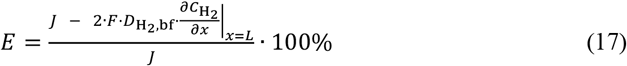

### Maximum current density

The MES-BIOFILM model was also used to determine which current density results in H_2_ saturation at the cathode surface, i.e. what is the maximum current density that can be applied to the cathode before H_2_ gas bubbles start to form and disrupt the biofilm. To find the maximum applicable current *J*_max,bf_, the system of ODEs was solved as described above with *J* as additional variable and with an additional constraint that saturation is reached at the cathode surface:

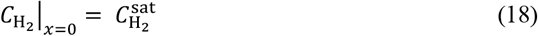

In Python, *J*_max,bf_ was obtained by minimizing the difference between 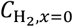 and 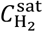using an error function.

### Maximum current density for bare cathode

To compare the case of a biofilm with an uncovered (bare) cathode, the maximum current density before H_2_ bubble formation starts (i.e. current density resulting in H_2_ saturation at the cathode surface) was also calculated for a bare electrode. In the case of a bare cathode, the flux of H_2_ generated by the cathode must equal the convective flux of H_2_ towards the bulk liquid:

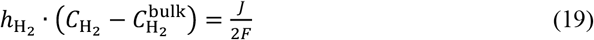

In which 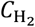 is the dissolved H_2_ concentration on the bare cathode surface. The maximum current density for an uncovered cathode *J*_max,bare_ is found when H_2_ saturation is reached on the cathode surface, or:

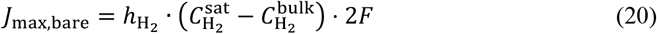

To quantify the effect of a biofilm in comparison to a bare electrode, a biofilm enhancement factor (EF) is defined as:

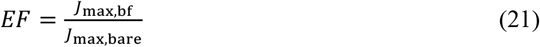

### Biofilm thickness at which DIC limitations arise

The thickness of the biofilm at which the DIC concentration becomes limiting was identified by determining the thickness for which the DIC concentration at the cathode-biofilm reaches zero. Above this critical thickness, a portion of the biofilm in proximity of the cathode would become inactive due to the local depletion of DIC and would no longer contribute to acetate production, resulting in an underutilization of the biofilm.

The limiting condition was identified numerically using a binary search over a predefined range of biofilm thicknesses. At each iteration, the model was evaluated at the middle point of the thickness range, and the resulting concentrations profiles were assessed for their physical feasibility, primarily verifying that the DIC concentration remained positive throughout the entire biofilm. A positive concentration of H_2_ as well as the expected monotonic trends for both concentration profiles were also considered. If the solution was physically feasible (implying

*C*_DIC_|_*x*=0_ > 0), the midpoint was taken as the new lower-bound of the biofilm thickness interval, while the upper bound was kept unchanged. On the other hand, if the solution was not feasible, the midpoint became the new upper bound, while the lower bound remained unchanged. This procedure progressively narrowed down the search interval until a prescribed precision of 10 µm was reached.

### Experimental measurement of the binary H_2_ biofilm diffusion coefficient

To estimate the binary diffusion coefficient of H_2_ in a (synthetic) biofilm, H_2_ concentration profiles were measured in an agar layer. The 2.5% agar layer was prepared by adding 25 g/L agar to 200 mL autotrophic growth medium for acetogenic bacteria (Grøn *et al*., 2026). This suspension was autoclaved in a serum bottle closed with a rubber stopper and next cooled down on a level plane to create a horizontally flat gel surface. The thickness of the 2.5% agar layer was 5.5 cm. After solidification, the headspace of the serum bottle was flushed with H_2_:CO_2_ (2:1) gas for 30 minutes and placed at 30 °C for 28 hours. A microsensor setup (Sandfeld *et al*., 2024) was used to profile the dissolved H_2_ concentration in the gel layer.

The binary H_2_ diffusion coefficient was obtained by fitting the experimental data with the analytical solution for transient diffusion in a slab with a fixed concentration as top boundary condition and zero flux as bottom boundary condition (Supplementary Method 2).

The obtained H_2_ diffusion coefficient for the gel (Table 1) was about 47% of the diffusion coefficient of H_2_ in water (Table 1). This ratio matches well with previously reported ratios of diffusion coefficients in natural biofilms versus in water (ranges between 0.2 and 0.8, with a mean of 0.4) (Stewart, 2003). The value for the biofilm diffusion coefficient calculated here can thus be considered representative for both synthetic and natural biofilms. In addition, we assumed that the biofilm reduced the CO_2_ diffusion coefficient with the same proportion as the H_2_ diffusion coefficient in comparison to water.

### Mass transfer coefficients

Realistic H_2_ and DIC mass transfer coefficients for the convective transport between the bulk liquid and the biofilm (Table 1) were calculated for a diffusive boundary layer on the cathode with a thickness of 200 µm (Rivera-Alvarez *et al*., 2020). The convective mass transfer at the right-hand boundary of the biofilm (Figure 1) can indeed be considered as diffusive mass transfer through a diffusive boundary layer adjacent to the biofilm or bare cathode. The convective mass transfer coefficient can therefore be calculated from the thickness of the diffusive boundary layer according to (assuming the Sherwood number equals 1) (Bergman *et al*., 2017):

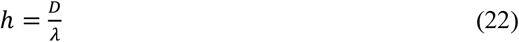

Where *h* is the mass transfer coefficient, *D* is the diffusion coefficient in the bulk liquid and *λ* is the thickness of the diffusive boundary layer.

### Relevant biofilm cell densities

A range of relevant biofilm cell densities (Table 1) was selected based on previously reported biofilm densities (Table S1). In general, biomass densities in microbial biofilms range from 0.3 up to 400 mg dry weight (DW) biomass per·cm^3^ biofilm (Table S1), while the fraction of active biomass (active cells) likely ranges between 10 and 90%, as a significant part of the biomass can consist of dead cells and the extracellular matrix (Flemming & Wingender, 2010, Rittmann & McCarty, 2020). The physical limit to the biofilm density is about 400 mg DW cm^-3^ biofilm, as this represents a biofilm completely filled with cells, as bacterial cell densities are about 400 mg cell DW per cm^-3^ cell volume (Table S1). Interestingly, the highest reported biofilm density during microbial electrosynthesis is 400 mg DW cm^-3^ biofilm (Table S1) and thus the theoretical maximum. For these reasons, we used a biofilm cell density of 400 mg DW·cm^-3^ (also gDW per L) biofilm as upper limit, while we used 100 mg active biomass per·cm^3^ biofilm as base case scenario.

### Other parameters

All parameter values for MES-BIOFILM are given in Table 1. The values of the different variables used to generate the figures are given in Table S2.

## Results

In this study, the model MES-BIOFILM was developed to investigate H_2_ and DIC concentration profiles in cathodic biofilms. For a first set of scenarios, we assumed that DIC was not limiting microbial H_2_ consumption. This is most likely the case in relatively thin biofilms, as the saturated H_2_ concentration is more than 200 times lower than the saturated DIC concentration (Table 1), while stoichiometrically double as much H_2_ than CO_2_ is consumed (Reaction 1). Moreover, we assumed that only very low DIC concentrations limit microbial consumption by using Monod kinetics with low *K*_m_ (Table 1). In this work, we define biofilms up to 200 µm as thin. We report below the specific biofilm thicknesses at which our model predicts DIC limitation.

### H_2_ concentration profiles in thin biofilms

Our simulations of the H_2_ concentration profiles in thin cathodic biofilms show that the dissolved H_2_ concentration exponentially decreases with increasing distance from the cathode surface (Figure 2). This decrease is the result of the combination of diffusion towards the bulk liquid and H_2_ consumption by the acetogens in the biofilm. The concentration profiles also show that the H_2_ concentration at the cathode surface (left-hand boundary, 0 µm) increases when higher current densities are applied (Figure 2A). Moreover, at high current densities (starting at ∼1.0 mA/cm^2^ for conditions used in our simulation), the H_2_ concentration at the cathode surface exceeds H_2_ saturation, meaning H_2_ bubbles would form and disrupt the biofilm, turning our model simulations unfeasible.

**Figure 2:**
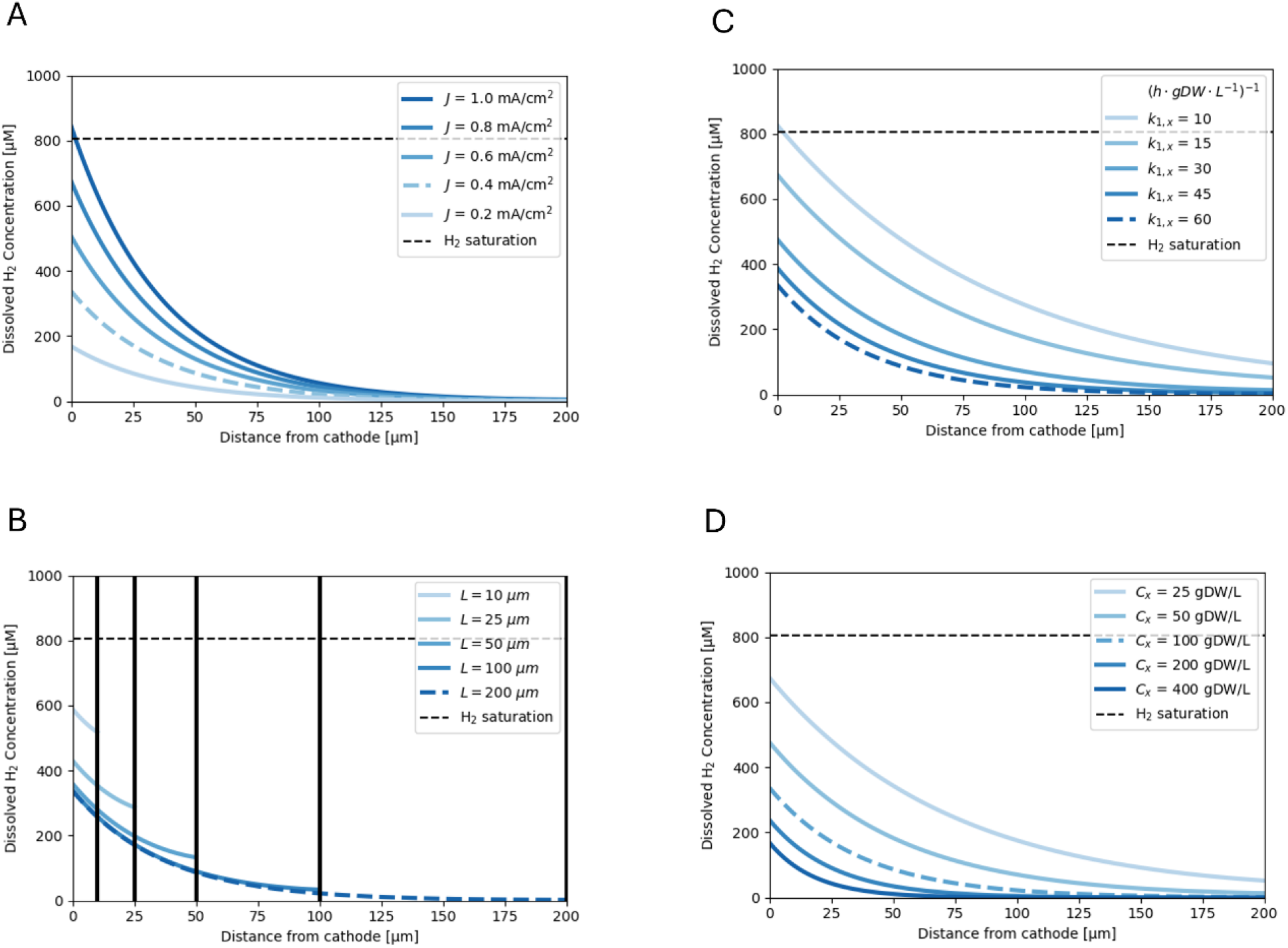
A: Simulated H_2_ concentration profiles in thin cathodic biofilms (up to 200 µm) with (A) different applied current densities, (B) different biofilm thicknesses, (C) different first-order rate coefficients and (D) different biofilm cell densities. Each plot changes only one variable, the blue long-dashed line represents the same simulation (identical conditions) in each panel. All other parameter values are given in Table S2. The black short-dashed line indicates H_2_ saturation. Any H_2_ concentration profile exceeding H_2_ saturation would signify *H*_*2*_ bubble formation, turning that simulation invalid.

Figure 2B shows how the H_2_ concentration profiles depend on the thickness of the biofilm. In very thin biofilms (10-50 µm), the H_2_ concentration at the outer edge of the biofilm (right-hand boundary) does not reach zero. This means that the microbial activity in these thin biofilms is insufficient to consume all H_2_ within the biofilm and that a large part of the generated H_2_ diffuses out of the biofilm into the bulk liquid. In contrast, in thicker biofilms, most of the H_2_ already gets consumed within the first 100 µm, while practically no H_2_ diffuses out of these biofilms (seen as the flat concentration gradient at the right-hand boundary) (Figure 2B). The effect of different parameters on the H_2_ utilization efficiency, i.e. fraction of cathodically generated H_2_ converted in the biofilm, is further discussed below (Figure 3). Figure 2B also shows that thicker biofilms lead to lower H_2_ concentrations throughout the biofilm at the same applied current density. Thicker biofilms thus also lower the H_2_ concentration at the cathode surface (left-hand boundary), thanks to their larger microbial activity.

**Figure 3:**
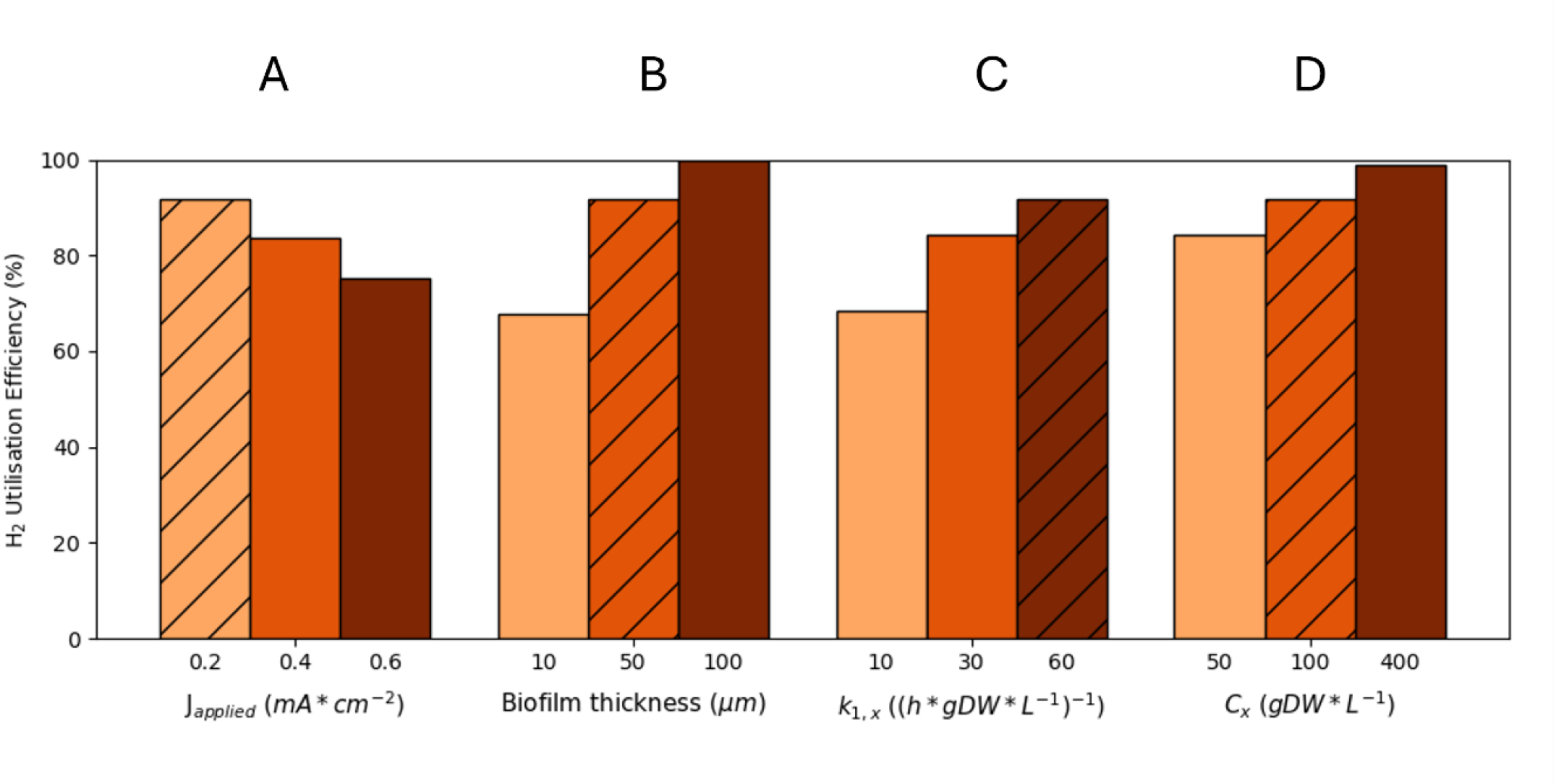
Simulated H_2_ utilization efficiencies in very thin cathodic biofilms (50 µm, unless stated differently) with (A) different applied current densities, (B) different biofilm thicknesses, (C) different first-order rate coefficients and (D) different biofilm cell densities. Each plot changes only one variable, other parameter values can be found in Table S2. The strikethrough bar represents the same simulation (identical conditions) in each panel.

At a fixed biofilm thickness, the H_2_ concentration profiles also depend on the microbial parameters for H_2_ consumption (Figure 2C and D). A higher biomass-specific first order rate coefficient 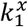was found to lower the H_2_ concentration profiles and the H_2_ concentration at the cathode surface due to the faster H_2_ consumption (Figure 2C). Previously, we reported that 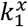 values differ up to six times among a range of acetogens, with the lowest 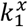 for *Sporomusa sphaeroides* (10.63 h^−1^×(gDW·L^−1^)) and the highest for *Acetobacterium wieringae* (60.93 h^−1^×(gDW·L^−1^)) (Munoz-Duarte *et al*., 2025). The H_2_ concentration profile in a biofilm thus depends on the specific acetogen and its H_2_ consumption characteristics. In addition, we previously reported that H_2_ thresholds vary up to three orders of magnitude among acetogens (Munoz & Philips, 2023), but the H_2_ threshold was found here to only have a minor impact on the H_2_ concentration profiles in cathodic biofilms (Figure S2). In contrast, the biofilm cell density affects the H_2_ concentration profiles in a similar manner as 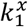, as a higher cell densities in the biofilm results in faster H_2_ consumption and thus also lowers the H_2_ concentration profile and the H_2_ concentration at the cathode surface e (Figure 2D).

### H_2_ utilization efficiency in 50 µm biofilms

Figure 2 suggested that in most 200 µm biofilms, all H_2_ gets fully consumed within the biofilm, which thus corresponds to 100% H_2_ utilization efficiency. In contrast, H_2_ does not get fully consumed in very thin biofilms (Figure 2B), leading to the loss of H_2_ to the bulk liquid and thus a lower H_2_ utilization efficiency. We therefore calculated the effect of different variables on the H_2_ utilization efficiency in 50 µm biofilms (Figure 3). The H_2_ utilization efficiency strongly depends on the applied current, as higher currents lower the H_2_ utilization efficiency (Figure 3A). At the same applied current, thinner biofilms lead to lower H_2_ utilization efficiencies (Figure 3B), as already described above. Nevertheless, 50 µm biofilms could still have a high H_2_ utilization efficiency upon increasing 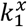 (Figure 3C) and the biofilm cell density (Figure 3D). Overall, our simulations suggest that very thin biofilms (50 µm) are already sufficient to convert most H_2_ resulting from a 0.2 mA·cm^-2^ current density, as long as the biofilm cell density and 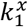 are maximized (Figure 3D, 400 g_DW_/L). Biofilms thinner than 50 µm cannot fully consume H_2_ within the biofilm, unless the applied current density is lowered.

### Maximum current densities and biofilm enhancement factors in thin biofilms

We calculated the maximum current density that can be applied before H_2_ gas bubbles are formed (Figure 4A and B), as the current density at which the H_2_ concentration at the cathode surface reaches saturation (Equation 18). This maximum current density increases with the cell density and 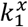 (Figure 4A). Our simulations suggest that in the most optimal conditions (highest 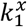 and highest biofilm cell density) a current density close to 2.0 mA/cm^2^ could be reached without H_2_ bubble formation on the cathode (Figure 4A). The maximum current density also increases with an increase of the biofilm thickness (Figure 4B), but only until a thickness of about 50 µm. Microbial activity beyond 50 µm likely does not affect the H_2_ concentration at the cathode surface, as is also suggested by Figure 2B. This thus means that very thin biofilms (∼50 µm) are already sufficient to suppress H_2_ bubble formation.

**Figure 4:**
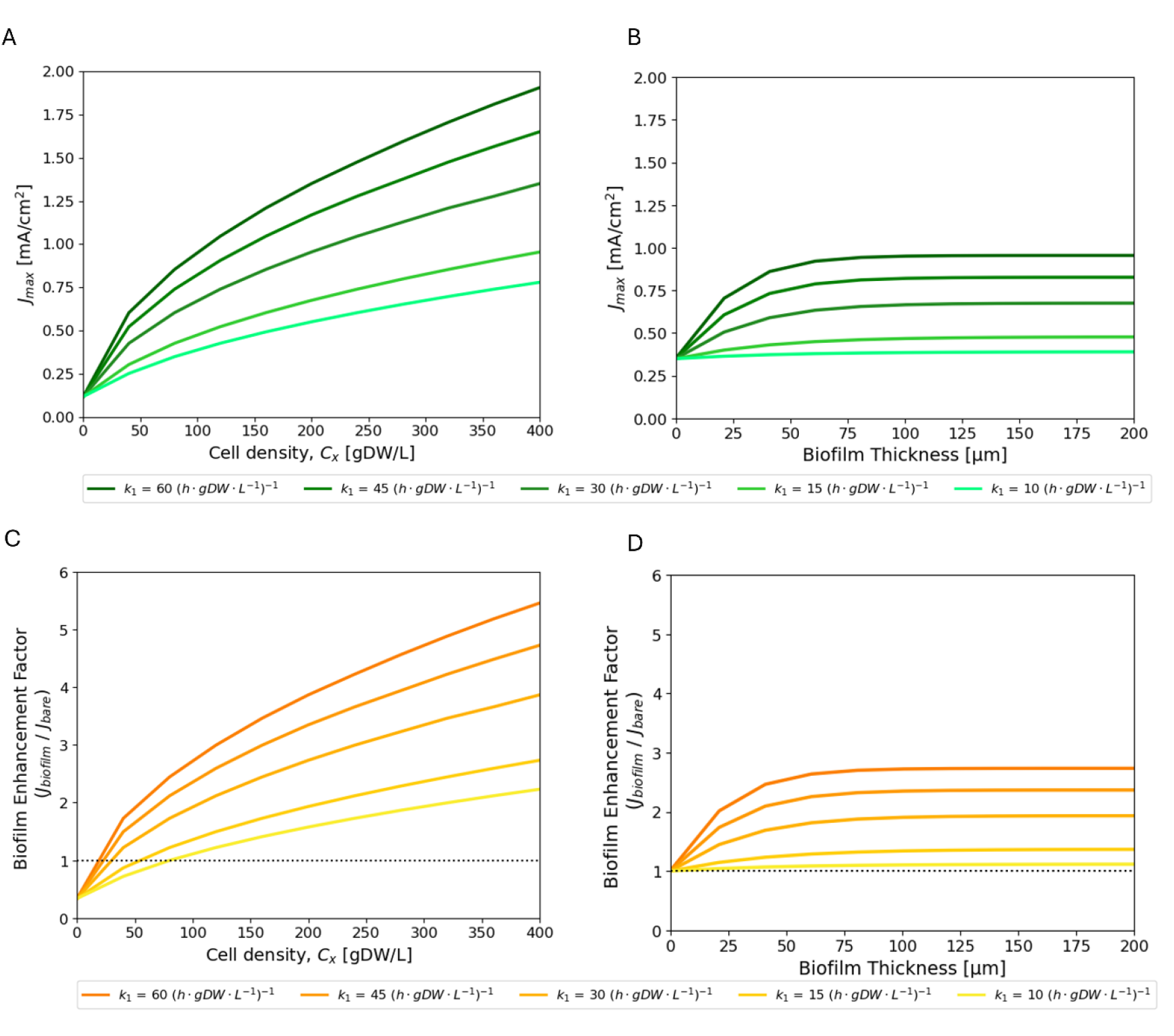
(A, B) Simulated maximum current density (i.e. highest current density that can be applied without exceeding H_2_ saturation) and (C, D) biofilm enhancement factors (i.e. factorial increase of the maximum current density due to a biofilm in comparison to a bare electrode) in function in function of (A,C) the biofilm cell density and (B, D) the biofilm thickness, as well as the first-order rate coefficient (A, B, C and D). Each plot changes two variables, other parameter values can be found in Table S2. Note that in B the case of a bare electrode is represented by the point where the simulations intercept with the Y-axis. Note that in C and D the dotted line represents a bare electrode.

In our conditions, the current density at which H_2_ bubbles start to form on a bare (uncovered) cathode (*J*_max,bare_) is 0.35 mA/cm^2^ (Equation 20). We further calculated biofilm enhancement factors (Figure 4C and D) by dividing the maximum current densities of cathodic biofilms with that of an uncovered electrode (Equation 21). For cathodic biofilms with a low cell density (<20-80 gDW/L), we found biofilm enhancement factors lower than 1, as in this case the biofilm rather acts as an insulating layer that prevents H_2_ transport to the bulk liquid. Such biofilms thus rather promote H_2_ bubble formation. In contrast, biofilms with a sufficiently high cell density resulted in biofilm enhancement factors higher than 1, illustrating their H_2_ bubble suppression effect. As for the maximum current density, the biofilm enhancement factor increases with the cell density and 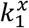 (Figure 4C) and with the biofilm thickness until ∼50 µm (Figure 4D). Furthermore, our simulations suggest that in the most optimal conditions (highest 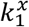 and highest biofilm cell density) a biofilm enhancement factor of 5.5 can be attained. This means that a 5.5 times higher current density can be applied to such a biofilm before H_2_ bubble formation starts in comparison to a bare electrode. Equation 20 shows that the maximum current density at a bare electrode depends on the H_2_ mass transfer coefficient *h*_H2_. This parameter was in this study kept constant, but any variations in 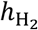 would thus also affect the biofilm enhancement factors.

### CO_2_ concentration profiles in thin biofilms

Next, we also simulated DIC concentration profiles in thin cathodic biofilms (200 µm) exposed to the maximum current densities described above. These simulations suggest that DIC concentrations remained high throughout the biofilms and thus did not limit H_2_ consumption (Figure 5). This is mainly because the DIC concentration in the bulk (set as the DIC concentration at pH 7 in equilibrium with CO_2_ partial pressure of 1 atm) is more than 200 times higher than the saturated H_2_ concentration (Table 1), while stoichiometrically double as much H_2_ than CO_2_ is consumed (Reaction 1).

**Figure 5:**
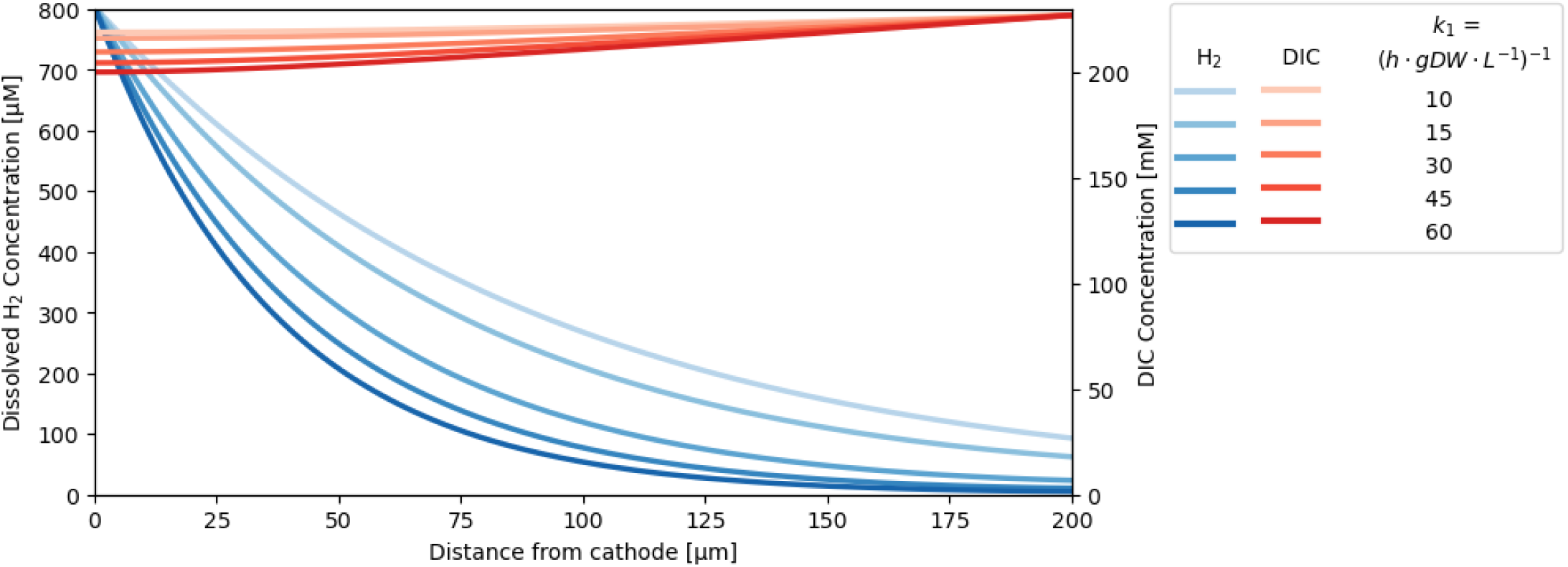
Simulated H_2_ and DIC concentration profiles at the maximum current density in cathodic biofilm with a thickness if 200 µm. This plot changes the first-order rate coefficient 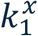, all other parameter values are given in Table S2. For each 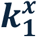, the H_2_ concentration profile at its maximum current density (Figure 4A) was first calculated. From this profile, the corresponding DIC concentration profile was simulated through conversion by stoichiometry.

### Biofilm thickness at which DIC limitations arise

Finally, we calculated at which biofilm thicknesses DIC limitations arise. For all the conditions included in our simulations, DIC limitations only started in biofilms thicker than 200 µm (Figure 6). All our simulations above, neglecting possible DIC limitations in thin biofilms (upto 200 µm), are thus valid. We further found that DIC limitations arise at thinner biofilms when microbial H_2_ and DIC consumption are higher (i.e. higher biofilm cell density and higher 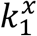) (Figure 6). In thick biofilms (1 mm, as in synthetic biofilms), DIC limitations are expected in all simulated cases, except when the biofilm cell density remains below 100 g_DW_/L in combination with slow H_2_ consumption (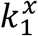 of 10 h^-1^·g_DW_^-1^·L).

**Figure 6:**
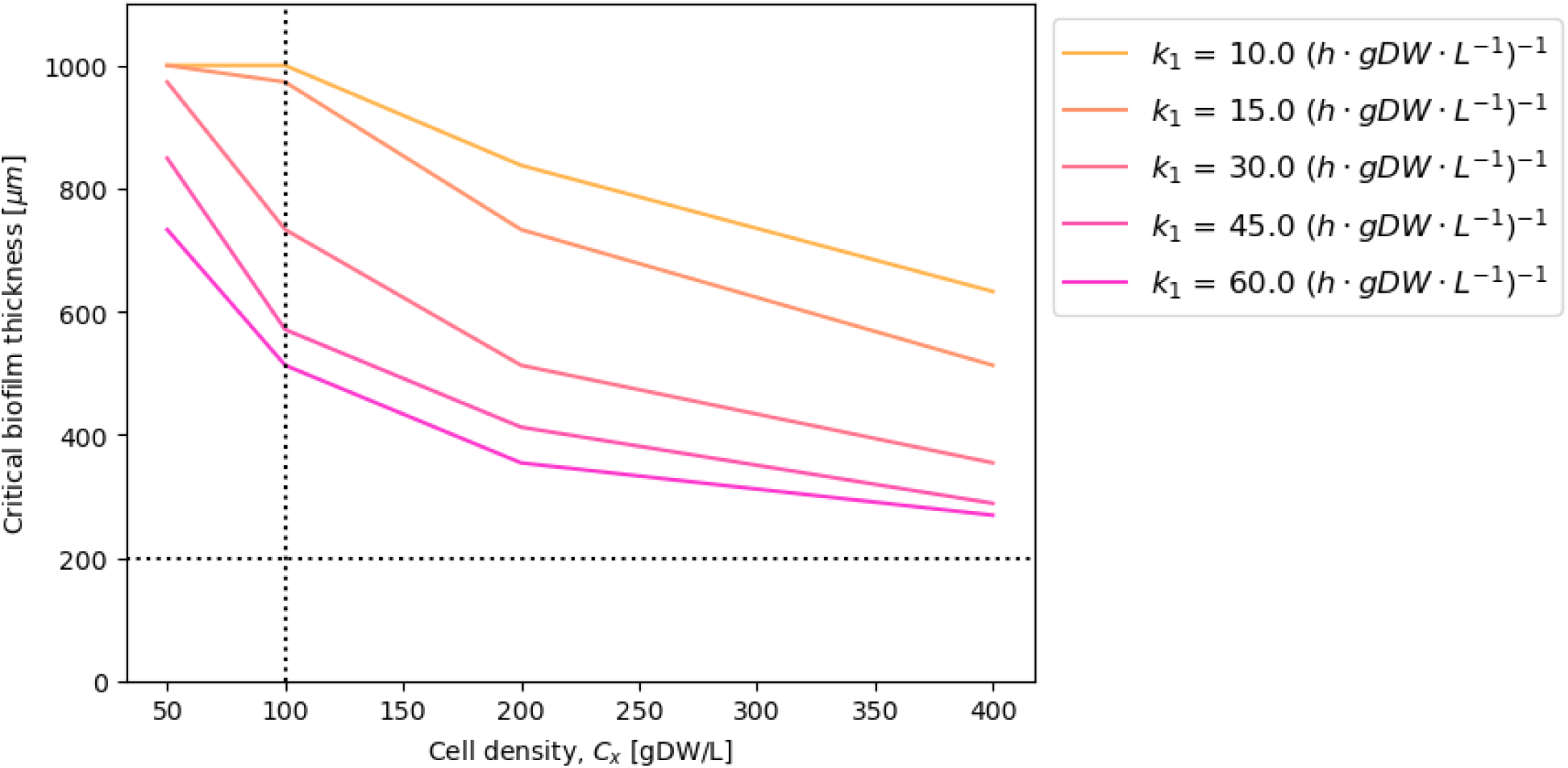
Critical biofilm thickness (i.e. biofilm thickness at which DIC limitations arise) as a function of the biofilm cell density and the first-order rate coefficient. This plot changes the biofilm cell density and 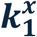; all other parameter values can be found in Table S2. Note that dotted lines represent a biofilm thickness of 200 µm and a biofilm cell density of 100 g_DW_/L, which are the base cases for the simulations above.

## DISCUSSION

### Implications for microbial electrosynthesis

In this work, we developed MES-BIOFILM, a comprehensive 1-D model, evaluating the importance of cathodic biofilms for microbial electrosynthesis systems. Our model simulations showed that cathodic biofilms can be effective in converting H_2_ and avoiding H_2_ transfer to the liquid phase (Figure 3). In addition, our results suggest that cathodic biofilms can suppress H_2_ bubble formation, as H_2_ consumption by the biofilm can keep the local H_2_ concentration under saturation (Figure 2, Figure 4). Interestingly, we calculated that this bubble suppression enables up to 5.5 times higher current densities in comparison to a bare electrode (Figure 4C).

In general, our study found that the contribution of a cathodic biofilm strongly depends on (1) the first order-rate coefficient, (2) the biofilm cell density; and (3) the biofilm thickness (Figure 2-4). The first order-rate coefficient can likely be controlled by selecting a pure culture with high reported *k*_*1*_^*x*^ value, e.g. *Acetobacterium wieringae* or *Sporomusa ovata* strain 2663 (Munoz-Duarte *et al*., 2025). In contrast, it is impossible to control and even accurately determine biofilm cell densities of cathodic biofilms. Biomass densities up to the theoretical maximum have been reported for MES biofilms (Cabau-Peinado *et al*., 2024) (Table S1), but this biofilms possibly contained a large inactive fraction, not contributing to H_2_ conversion. Nevertheless, very dense and active biofilms are required to maximize the effect of a cathodic biofilm for MES, as suggested by our simulations (Figure 4).

MES-BIOFILM showed that biofilms should be at least 50 µm thick to fully convert H_2_ within the biofilm (depending on the applied current density (Figure 3)) and to maximally suppress H_2_ bubble formation (Figure 4). However, cathodic biofilms, definitely those of pure culture acetogens, are mostly thinner (overview given in (Grøn *et al*., 2026)), while cathodic biofilms usually grow very slow (Vassilev *et al*., 2022). Synthetic biofilms could overcome this slow biofilm growth and thin natural biofilms (Krige *et al*., 2021) and possibly allow to control the biofilm thickness and cell density. However, our model simulations suggest that in thick biofilms (the synthetic biofilms of Krige et al. (2021) were 1 mm), DIC diffusion limitations arise in almost all conditions (Figure 5). As biofilm parameters, i.e. biofilm thickness and cell density, are so far hard to control, it will be hard to use cathodic biofilms in practice to increase the efficiency of MES and suppress H_2_ bubble formation. Even an experimental validation of our model results will be hard without experimental control of biofilm parameters.

MES-BIOFILM suggests that an optimal cathodic biofilm can handle 5.5 times higher current densities in comparison to a bare electrode before H_2_ bubbles arise (Fig 4C). This offers interesting perspectives to increase current densities for MES without losing efficiency. Previously, several studies stated that cathodic biofilms rather limit the attainable current densities in MES (Claassens *et al*., 2019, Wood *et al*., 2021), but these studies did not consider the effect of biofilms on H_2_ bubble suppression. Our simulations showed that sparse biofilms (low cell density) do have a negative effect on bubble suppression (lower maximum current density than with bare electrode) (Figure 4C), but that biofilms with a high cell density have a strong bubble suppression effect. Nevertheless, it is questionable if a 5.5-times current density enhancement is sufficient to bring MES closer to commercial relevance. Prévoteau et al. (2020) described that current densities in MES should at least increase 30 times to reach those of commercial electrolysers. We consider that mainly the theoretical maximum biofilm cell density (Table S1) restricts a further increase of the biofilm enhancement factor. One feasible option to further increase the biofilm effect might be strain improvement to increase the H_2_ consumption rates (effect of *k*_1_^x^ in Figure 4). For this reason, cathodic biofilms alone will likely not be enough to optimize MES and will need to be combined with a high electrode surface area and proper reactor design to increase the overall current and rate of MES.

### Study limitations

It is important to note that MES-BIOFILM did not incorporate the effect of the acetate concentrations nor pH on the microbial H_2_ consumption rate. Nevertheless, deviations in both could affect acetogen H_2_ consumption (LaBelle *et al*., 2014, Ntagia *et al*., 2020, Kwon *et al*., 2022). Some scientists have even stated that proton diffusion towards the cathode could limits electrochemical H_2_ generation, thereby creating strong pH gradients (Kracke *et al*., 2021). Strong pH gradients, however, can likely be minimized by sufficient pH buffering. Moreover, protons do not only migrate due to diffusion, but also by the electric field attracting the positively charged ions towards the cathode (Marcus *et al*., 2010). For these reasons, we expect that strong pH gradients in cathodic biofilms only arise at current densities higher than those onsetting H_2_ bubble formation, which were out of scope of this study. We thus assume it was fair to neglect proton transport as a limiting factor in our study. Nevertheless, our biofilm enhancement factors (Fig 4 and C) should best be considered as maximum attainable values.

MES-BIOFILM did incorporate the effect of the DIC concentration on the microbial H_2_ consumption rate (Equation 4). However, our model used kinetic parameters for methanogens (Table 1), as so far, no study has investigated CO_2_ consumption kinetics of acetogens. Previously, we found that the H_2_ consumption kinetics of acetogens are strongly different from those of methanogens (first order vs Monod kinetics) (Munoz-Duarte *et al*., 2025), emphasizing the need for accurate kinetic parameters to further increase the predictive power of model simulations.

Our study assumed that H_2_ bubble formation starts once H_2_ saturation is reached. In practice, local oversaturation is often required before bubble nucleation occurs (Carvajalino-Olave *et al*., 2025). The H_2_ concentration at which bubble nucleation starts, depends on various factors including the liquid-gas surface tension and the electrode surface roughness and chemistry (Carvajalino-Olave *et al*., 2025). The possible effect of these physical properties on the microbial H_2_ bubble suppression warrants further investigations.

Furthermore, MES-BIOFILM assumed a flat and homogenous cathode material for reasons of simplicity. In practice, cathodes for MES often consist of 3D porous materials. Such electrode materials enhance the attainable current densities (Jourdin & Burdyny, 2021), but probably also lead to heterogenic H_2_ bubble formation, as H_2_ bubble nucleation preferably occurs at electrode edges. Future modelling studies should therefore also incorporate the complexity of porous electrode structures to further evaluate the importance of microbial H_2_ bubble suppression in more realistic systems.

### Overall conclusion

Overall, our model simulations suggest that cathodic biofilms can play an important role in microbial electrosynthesis, as long as they are sufficiently thick and dense in cells. The first studies on MES assumed a dependency on direct extracellular electron uptake (Cheng *et al*., 2009, Nevin *et al*., 2010, Nevin *et al*., 2011, Kato *et al*., 2015), for which a biofilm is obviously essential. However, as the important role of H_2_ as key intermediate in MES through indirect extracellular electron uptake is now recognized (Blanchet *et al*., 2015, Philips, 2020, Boto *et al*., 2023), the importance of cathodic biofilms has become less clear. Several studies have even argued against cathodic biofilms, as they would limit the current density and available rates (Claassens *et al*., 2019, Wood *et al*., 2021), while others have nevertheless observed biofilms in MES system with a high production (LaBelle & May, 2017, Cabau-Peinado *et al*., 2024). Our modelling results suggest that cathodic biofilms can play an important role in both (1) the efficient conversion of H_2_ (no loss of H_2_ to the liquid and ultimately to gas phase) and (2) in suppression the onset of H_2_ bubble formation (allowing applying higher current densities). However, cathodic biofilms alone are likely insufficient to bring MES to the commercially relevant high current densities. So far, these insights are only based on modelling simulations and will still need further investigation by experimental studies.

## ACKNOWLEDGEMENTS

This research was funded by a Green Transition Project I grant (MES-MODEL, 1127-00048B) from the Independent Research Fund Denmark (DFF), in addition to a grant for the Novo Nordisk Foundation CO_2_ Research Center (NNF21SA0072700). We thank Petra Fangel Dürr and Louise Vinther Grøn to their contribution to determine the biofilm diffusion coefficient.

## AUTHOR CONTRIBUTIONS

Mikkel Bendix Jensen and Jacopo Catalano designed and developed the model. Vladimir Koselka ran the simulations and performed the visualisations. Jo Philips wrote the manuscript, while all other authors edited the manuscript. Jo Philips and Jacopo Catalano acquired funding and performed supervision. All authors contributed to the article and approved the submitted version.

## REFERENCES

Bergman TL, Lavine AS, Incropera FP & DeWitt DP (2017) Fundamentals of Heat and Mass Transfer. Wiley.

Blanchet E, Duquenne F, Rafrafi Y, Etcheverry L, Erable B & Bergel A (2015) Importance of the hydrogen route in up-scaling electrosynthesis for microbial CO2 reduction. Energy & Environmental Science 8: 3731–3744.

Boto ST, Bardl B, Harnisch F & Rosenbaum MA (2023) Microbial electrosynthesis with Clostridium ljungdahlii benefits from hydrogen electron mediation and permits a greater variety of products. Green Chemistry 25: 4375–4386.

Braus-Stromeyer SA, Schnappauf G, Braus GH, Gössner AS & Drake HL (1997) Carbonic anhydrase in Acetobacterium woodii and other acetogenic bacteria. Journal of Bacteriology 179: 7197–7200.

Cabau-Peinado O, Straathof AJJ & Jourdin L (2021) A general model for biofilm-driven microbial electrosynthesis of carboxylates from CO2. Frontiers in Microbiology Volume 12 – 2021.

Cabau-Peinado O, Winkelhorst M, Stroek R, de Kat Angelino R, Straathof AJJ, Masania K, Daran JM & Jourdin L (2024) Microbial electrosynthesis from CO2 reaches productivity of syngas and chain elongation fermentations. Trends in Biotechnology 42: 1503–1522.

Callander G, Deutzmann J & Spormann A (2024) Alkaline hydrogenotrophic methanogenesis in Methanococcus vannielii at low carbon dioxide concentrations. Journal of CO2 Utilization 83: 102788.

Carvajalino-Olave J-D, Laguillaumie L, Dumas C & Erable B (2025) Electroreduction of water, a promising alternative to gas injection: A case on hydrogen solubilization in biological processes involving hydrogenotrophs. Chemical Engineering Journal 523: 168494.

Cheng S, Xing D, Call DF & Logan BE (2009) Direct biological conversion of electrical current into methane by electromethanogenesis. Environmental Science & Technology 43: 3953–3958.

Claassens NJ, Cotton CAR, Kopljar D & Bar-Even A (2019) Making quantitative sense of electromicrobial production. Nature Catalysis 2: 437–447.

Cussler EL (2009) Diffusion - Mass Transfer in Fluid Systems. Cambridge University Press.

Deutzmann JS & Spormann AM (2024) High acetate titer obtained from CO2 by thermophilic microbial electrosynthesis with Thermoanaerobacter kivui. Bioresource Technology Reports 25: 101740.

Drake HL, Gossner AS & Daniel SL (2008) Old acetogens, new light. Incredible Anaerobes: From Physiology to Genomics to Fuels, Vol. 1125 (Wiegel J, Maier RJ & Adams MWW, eds.), p.^pp. 100–128. Blackwell Publishing, Oxford.

Flemming H-C & Wingender J (2010) The biofilm matrix. Nature Reviews Microbiology 8: 623–633.

Grøn LV, Muñoz-Duarte L, Marshall IPG, Johnsen NK, Eser BE, Ma Y, Eken M, Kofoed MVW, Koren K & Philips J (2026) Adaptive laboratory evolution increased biofilm formation by Sporomusa ovata through a mutation in galU. Biofilm 11: 100368.

Jourdin L & Burdyny T (2021) Microbial electrosynthesis: Where do we go from here? Trends Biotechnol 39: 359–369.

Kato S, Yumoto I & Kamagata Y (2015) Isolation of acetogenic bacteria that induce biocorrosion by utilizing metallic iron as the sole electron donor. Applied and Environmental Microbiology 81: 67–73.

Kazemi M, Biria D & Rismani-Yazdi H (2015) Modelling bio-electrosynthesis in a reverse microbial fuel cell to produce acetate from CO2 and H2O. Physical Chemistry Chemical Physics 17: 12561–12574.

Kolev NI (2011) Multiphase Flow Dynamics 4: Turbulence, Gas Adsorption and Release, Diesel Fuel Properties. Springer.

Kracke F, Deutzmann JS, Jayathilake BS, Pang SH, Chandrasekaran S, Baker SE & Spormann AM (2021) Efficient hydrogen delivery for microbial electrosynthesis via 3D-printed cathodes. Frontiers in Microbiology Volume 12 – 2021.

Krige A, Rova U & Christakopoulos P (2021) 3D bioprinting on cathodes in microbial electrosynthesis for increased acetate production rate using Sporomusa ovata. Environmental Chemical Engineering 9: 106189–106189.

Kuchenbuch A, Al-Sbei S, Rosa LFM, Boto ST, Westermann M, Rosenbaum MA & Harnisch F (2025) Reducing oxygen stress and improving hydrogen availability boosts microbial electrosynthesis by Clostridium ljungdahlii. ChemSusChem 18: e202501118.

Kwon SJ, Lee J & Lee HS (2022) Metabolic changes of the acetogen Clostridium sp. AWRP through adaptation to acetate challenge. Frontiers in Microbiology Volume 13 – 2022.

LaBelle EV & May HD (2017) Energy efficiency and productivity enhancement of microbial electrosynthesis of acetate. Frontiers in Microbiology Volume 8 – 2017.

LaBelle EV, Marshall CW, Gilbert JA & May HD (2014) Influence of acidic pH on hydrogen and acetate production by an electrosynthetic microbiome. PLOS ONE 9: e109935.

Li Z, Fu Q, Su H, Yang W, Chen H, Zhang B, Hua L & Xu Q (2022) Model development of bioelectrochemical systems: A critical review from the perspective of physiochemical principles and mathematical methods. Water Research 226: 119311.

Marcus AK, Torres CI & Rittmann BE (2010) Evaluating the impacts of migration in the biofilm anode using the model PCBIOFILM. Electrochimica Acta 55: 6964–6972.

Munoz-Duarte L, Chakraborty S, Gron LV, Bambace MF, Catalano J & Philips J (2025) H2 consumption by various acetogenic bacteria follows first-order kinetics up to H2 saturation. Biotechnol Bioeng 122: 804–816.

Munoz L & Philips J (2023) No acetogen is equal: Strongly different H(2) thresholds reflect diverse bioenergetics in acetogenic bacteria. Environ Microbiol 25: 2032–2040.

Murugesan V, Hartkamp R & Padding JT (2026) Microbial electrosynthesis for CO2 conversion: Process limiting steps investigated by micro-scale modeling. Electrochimica Acta 557: 148494.

Nevin KP, Woodard TL, Franks AE, Summers ZM & Lovley DR (2010) Microbial electrosynthesis: Feeding microbes electricity to convert carbon dioxide and water to multicarbon extracellular organic compounds. Mbio 1.

Nevin KP, Hensley SA, Franks AE, Summers ZM, Ou J, Woodard TL, Snoeyenbos-West OL & Lovley DR (2011) Electrosynthesis of organic compounds from carbon dioxide Is catalyzed by a diversity of acetogenic microorganisms. Applied and Environmental Microbiology 77: 2882–2886.

Newell DB & Tiesinga E (2019) The International System of Units (SI).

Ntagia E, Chatzigiannidou I, Williamson AJ, Arends JBA & Rabaey K (2020) Homoacetogenesis and microbial community composition are shaped by pH and total sulfide concentration. Microbial Biotechnology 13: 1026–1038.

Philips J (2020) Extracellular electron uptake by acetogenic bacteria: Does H2 consumption favor the H2 evolution reaction on a cathode or metallic iron? Frontiers in Microbiology 10: 2997.

Prevoteau A, Carvajal-Arroyo JM, Ganigue R & Rabaey K (2020) Microbial electrosynthesis from CO2: forever a promise? Curr Opin Biotechnol 62: 48–57.

Rabaey K & Rozendal RA (2010) Microbial electrosynthesis - revisiting the electrical route for microbial production. Nature Reviews Microbiology 8: 706–716.

Ragab Aa, Shaw DR, Katuri KP & Saikaly PE (2020) Effects of set cathode potentials on microbial electrosynthesis system performance and biocathode methanogen function at a metatranscriptional level. Scientific Reports 10: 19824.

Rittmann BE & McCarty PL (2020) Environmental Biotechnology: Principles and Applications. McGraw-Hill.

Rivera-Alvarez I, Brown RK, Keskin-Pyttel D, Steffens J, Farber P & Schröder U (2020) Correlating theoretical boundary layer thickness to the power output of a microbial fuel cell with a complex anode geometry operated at varying flow rates. Journal of Power Sources 470: 228428.

Rominger L, Hackbarth M, Jung T, Scherzinger M, Rosa LFM, Horn H, Kaltschmitt M, Picioreanu C & Gescher J (2025) Efficiency and process development for microbial biomass production using oxic bioelectrosynthesis. Trends in Biotechnology 43: 673–695.

Sandfeld T, Grøn LV, Munoz L, Meyer RL, Koren K & Philips J (2024) Considerations on the use of microsensors to profile dissolved H2 concentrations in microbial electrochemical reactors. PLoS ONE 19: e0293734–e0293734.

Stewart PS (2003) Diffusion in biofilms. Journal of Bacteriology 185: 1485–1491.

Vassilev I, Dessì P, Puig S & Kokko M (2022) Cathodic biofilms – A prerequisite for microbial electrosynthesis. Bioresource Technology 348: 126788–126788.

Wood JC, Grové J, Marcellin E, Heffernan JK, Hu S, Yuan Z & Virdis B (2021) Strategies to improve viability of a circular carbon bioeconomy - A techno-economic review of microbial electrosynthesis and gas fermentation. Water Research 201: 117306–117306.

