## supplementary information file for "1D model simulations suggest biofilms can suppress cathodic H_2_ bubble formation"

*Corresponding author

**Supplementary Method 1: Derivation of altered boundary condition**

We estimated the maximum DIC flux at the outer boundary, by considering the maximum applied current density in our simulations (Figure 4A) and the complete consumption of all $H_2$ generated at the cathode within the biofilm. The corresponding maximum DIC flux is:

$$J_{\mathrm{DIC,\ max}}\Big|_{x=L} = -\frac{J_{\mathrm{max,bf}}}{4 \cdot F} \quad \text{(S1)}$$

According to the boundary condition (Eq 10), this becomes:

$$h_{\mathrm{DIC}}\left(C_{\mathrm{DIC}}|_{x=L} - C_{\mathrm{DIC}}^{\mathrm{bulk}}\right) \pounds -\frac{J_{\mathrm{max,bf}}}{4 \cdot F} \quad \text{(S2)}$$

Using $h_{\mathrm{DIC}} = 9.6$ µm/s (Table 1) and $J_{\mathrm{max}} = 2\ \mathrm{mA/cm^2}$ (Figure 4A), we obtain:

$$C_{\mathrm{DIC}}|_{x=L} - C_{\mathrm{DIC}}^{\mathrm{bulk}} \pounds \frac{J_{\mathrm{max,bf}}}{4 \cdot F} \cdot \frac{1}{h_{\mathrm{DIC}}} \sim 5.4\ \frac{\mathrm{mol}}{\mathrm{m^3}} \quad \text{(S3)}$$

The difference between the DIC concentration at the outer boundary and in the bulk is thus maximally 5.4 mM for the conditions of our simulations. As the bulk DIC concentration is set to 227 mM (Table 1), the DIC concentration difference between the outer boundary and the bulk is thus maximally ~2.4% and can be considered negligible in the conditions simulated in this work. Therefore, we can replace Eq. (10) in the main text with a fixed concentration boundary condition:

$$C_{\mathrm{DIC}}|_{x=L} = C_{\mathrm{DIC}}^{\mathrm{bulk}} \quad \text{(S4)}$$

**Supplementary method 2: Experimental measurement of biofilm diffusion coefficient**

The $H_2$ diffusion coefficient in an agar gel layer was determined as an approximation for the $H_2$ diffusion coefficient in synthetic or natural biofilms. The top of a 55 mm agar (2.5%) layer was exposed to a 66% $H_2$ atmosphere for a duration of 28 hours. Afterwards, the $H_2$ concentration profile in this gel layer was measured using a $H_2$ microsensor (Unisense, Denmark) up to a distance of 23 mm from the bottom of the gel (Figure S1).

The experimentally measured concentration profile was fitted with the analytical solution for transient diffusion in a slab with a fixed concentration at the top and a no-flux boundary at the bottom (Crank, 1975):

$$\frac{C-C_0}{C_l-C_0} = 1 - \frac{4}{\pi} \sum_{n=0}^{50} \frac{(-1)^n}{2n+1} \cdot e^{\frac{-D(2n+1)^2\pi^2 t}{4l^2}} \cdot \cos\left(\frac{(2n+1)\pi x}{2l}\right) \quad \text{(S5)}$$

Where $D$ is the diffusion coefficient to be estimated [$m^2 \cdot s^{-1}$], $l$ is the length of the gel layer in [m], $t$ is the time in [s], $x$ is the position of the measurement within the gel in [m], $C_0$ is the initial $H_2$ concentration throughout the gel (0 μM), $C_l$ is the fixed concentration at the top of the gel (set to be 400 μM, based on the experimental measurements after 28 hours), and $C$ is the concentration at position $x$ and time $t$.

The difference between the experimental and modelled datapoints was minimised using a Nelder-Mead algorithm. The fitted biofilm $H_2$ diffusion coefficient had a value of $2.27 \cdot 10^{-9}$ $m^2 \cdot s^{-1}$.

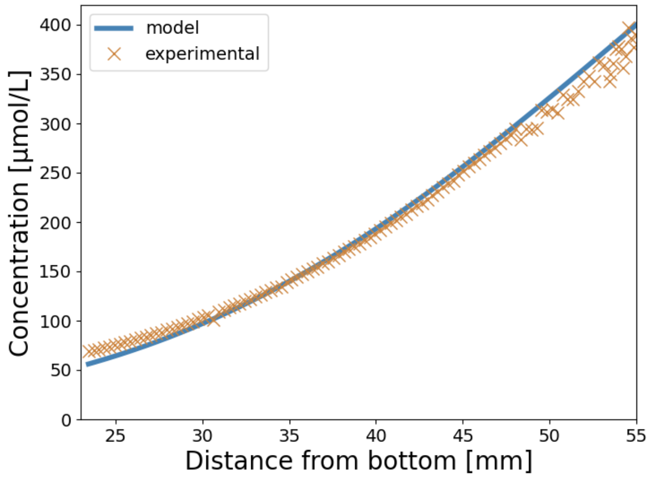


**Figure S1: Experimentally and fitted $H_2$ concentrations throughout the 55 mm agar layer that had been exposed for 28 hours to an atmosphere of 66% $H_2$.**

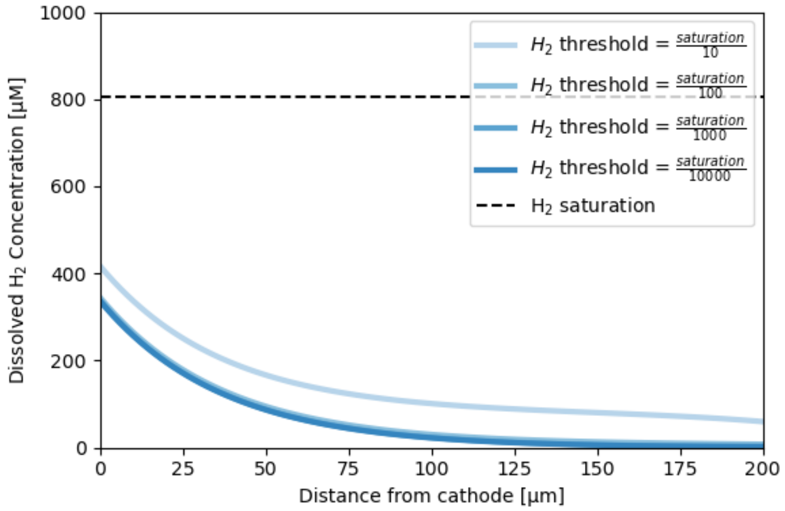


**Figure 12: Simulated $H_2$ concentration profiles in cathodic biofilms with $H_2$ thresholds. All other parameter values are given in Table S2 in the main text. The black short-dashed line indicates $H_2$ saturation.**

**Table S1: Different types of relevant biomass densities in dry weight (DW) per volume.**

| Biomass density (mg DW·cm$^{-3}$) | Description | Source |
|---|---|---|
| *Density of bacterial cells (cell dry weight per cell volume)* | | |
| 435 | Experimentally derived dry weight density of bacterial cells | (Loferer-Krößbacher *et al.*, 1998) |
| 330 | Converted from the reported wet cell density of 1.09 g·cm$^{-3}$ and average dry weight content of 30% for bacterial cells | (Bakken Lars & Olsen Rolf, 1983) |
| 300-400 | Experimental dry mass density for *Escherichia coli* and *Caulobacter crescentus* | (Oldewurtel *et al.*, 2021) |
| *Biomass density of microbial biofilms (biomass dry weight per biofilm volume)** | | |
| 0.3-115 | Converted from reported $10^8$-$10^{11}$ cells per g wet biofilm using correlation factors between cell numbers and dry weight for acetogens (Munoz-Duarte *et al.*, 2025), assuming 1 g wet biofilm corresponds with 1 cm$^3$. | (Flemming *et al.*, 2016) |
| 1-300 | Low biomass density expected for aerobic low-stress biofilms, much higher densities for anaerobic high-stress biofilms | (Rittmann & McCarty, 2020) |
| 25-48 | Biofilms grown on sucrose and $O_2$. Biomass density decreased with biofilm thickness. Biofilm thicknesses ranged from 100 to 1350 μm. | (Huang *et al.*, 1985) |
| 10-80 | Biofilms grown on sand particle in fluidized bed reactors fed with phenol. Less dense, when thicker biofilms. | (Ro & Neethling, 1991) |
| 100-200 | Experimental biofilm densities in porous cathodes during microbial electrosynthesis | (Winkelhorst *et al.*, 2023) |
| 400 | Maximum biofilm density in porous cathode during microbial electrosynthesis | (Cabau-Peinado *et al.*, 2024) |
| *Highest planktonic cell densities of acetogenic bacteria (dry cel weight per volume liquid phase)* | | |
| 10 | Maximum cell density of *Clostridium ljungdahlii* in gas fermentation system with cell retention | (Perret *et al.*, 2024) |
| 14 | Highest cell density of *Acetobacterium woodii* in gas fermentation system with cell retention | (Kantzow *et al.*, 2015) |

**The fraction active biomass (active cells) of the total biofilm biomass ranges between 10 and 90%, the remaining biomass consists of dead cells and the extracellular matrix (Flemming & Wingender, 2010, Rittmann & McCarty, 2020).*

**Table S2: Overview of parameter values used for the different figures.**

| Parameter | Biofilm thickness | First-order rate coefficient | $H_2$ threshold | Cell density | Mass transfer coefficient | Current density |
|---|---|---|---|---|---|---|
| Symbol | $L$ | $k_1^x$ | $C^{\star}_{H_2}$ | $C_X$ | $h_{H2}$ | $J$ |
| Unit | µm | $h^{-1} \cdot g_{DW}^{-1} \cdot L$ | µM | $g_{DW} \cdot L^{-1}$ | $\mu m \cdot s^{-1}$ | $mA \cdot cm^{-2}$ |
| Figure 2 | | | | | | |
| A | 200 | 60 | 0 | 100 | 22.5 | 0.2-1.0 |
| B | 10-200 | 60 | 0 | 100 | 22.5 | 0.4 |
| C | 200 | 10-60 | 0 | 100 | 22.5 | 0.4 |
| D | 200 | 60 | 0 | 25-400 | 22.5 | 0.4 |
| Figure 3 | | | | | | |
| A | 50 | 60 | 0 | 100 | 22.5 | 0.2-0.6 |
| B | 10-100 | 60 | 0 | 100 | 22.5 | 0.2 |
| C | 50 | 10-60 | 0 | 100 | 22.5 | 0.2 |
| D | 50 | 60 | 0 | 50-400 | 22.5 | 0.2 |
| Figure 4 | | | | | | |
| A | 200 | 10-60 | 0 | 0-400 | 22.5 | maximized |
| B | 0-200 | 10-60 | 0 | 100 | 22.5 | maximized |
| C | 200 | 10-60 | 0 | 0-400 | 22.5 | maximized |
| D | 0-200 | 10-60 | 0 | 100 | 22.5 | maximized |
| Figure 5 | | | | | | |
| | 200 | 10-60 | 0 | 100 | 22.5 | maximized |
| Figure 6 | Searched within | | | | | |
| | 0-1000 | 10-60 | 0 | 50-400 | 22.5 | maximized |
| Figure S2 | | | | | | |
| | 200 | 60 | 0.08-80 | 100 | 22.5 | 0.4 |

### REFERENCES


Bakken Lars R & Olsen Rolf A (1983) Buoyant densities and dry-matter contents of microorganisms: Conversion of a measured biovolume into biomass. *Applied and Environmental Microbiology* **45**: 1188-1195.

Cabau-Peinado O, Winkelhorst M, Stroek R, de Kat Angelino R, Straathof AJJ, Masania K, Daran JM & Jourdin L (2024) Microbial electrosynthesis from $CO_2$ reaches productivity of syngas and chain elongation fermentations. *Trends in Biotechnology* **42**: 1503-1522.

Crank J (1975) *The Mathematics of Diffusion*. Oxford Science Publications.

Flemming H-C & Wingender J (2010) The biofilm matrix. *Nature Reviews Microbiology* **8**: 623-633.

Flemming H-C, Wingender J, Szewzyk U, Steinberg P, Rice SA & Kjelleberg S (2016) Biofilms: An emergent form of bacterial life. *Nature Reviews Microbiology* **14**: 563-575.

Huang JC, Chang SY, Liu YC & Jiang Z (1985) Biofilm growths with sucrose as substrate. *Journal of Environmental Engineering* **111**: 353-363.

Kantzow C, Mayer A & Weuster-Botz D (2015) Continuous gas fermentation by *Acetobacterium woodi*i in a submerged membrane reactor with full cell retention. *Journal of Biotechnology* **212**: 11-18.

Loferer-Krößbacher M, Klima J & Psenner R (1998) Determination of bacterial cell dry mass by transmission electron microscopy and densitometric image analysis. *Applied and Environmental Microbiology* **64**: 688-694.

Munoz-Duarte L, Chakraborty S, Gron LV, Bambace MF, Catalano J & Philips J (2025) $H_2$ consumption by various acetogenic bacteria follows first-order kinetics up to $H_2$ saturation. *Biotechnol Bioeng* **122**: 804-816.

Oldewurtel ER, Kitahara Y & van Teeffelen S (2021) Robust surface-to-mass coupling and turgor-dependent cell width determine bacterial dry-mass density. *Proceedings of the National Academy of Sciences* **118**: e2021416118.

Perret L, Boukis N & Sauer J (2024) Synthesis gas fermentation at high cell density: How pH and hydrogen partial pressure affect productivity and product ratio in continuous fermentation. *Bioresource Technology* **391**.

Rittmann BE & McCarty PL (2020) *Environmental Biotechnology: Principles and Applications*. McGraw-Hill.

Ro KS & Neethling JB (1991) Biofilm density for biological fluidized beds. *Research Journal of the Water Pollution Control Federation* **63**: 815-818.

Winkelhorst M, Cabau-Peinado O, Straathof AJJ & Jourdin L (2023) Biomass-specific rates as key performance indicators: A nitrogen balancing method for biofilm-based electrochemical conversion. *Frontiers in Bioengineering and Biotechnology* **Volume 11 - 2023**.
